# Nuclear and plastid phylogenomic analyses provide insights into the reticulate evolution, species delimitation and biogeography of the Sino-Japanese disjunctive *Diabelia* (Caprifoliaceae)

**DOI:** 10.1101/2021.05.31.446416

**Authors:** Xiu-Rong Ke, Diego F. Morales-Briones, Hong-Xin Wang, Qing-Hui Sun, Jacob B. Landis, Jun Wen, Hua-Feng Wang

**Affiliations:** Key Laboratory of Tropical Biological Resources of Ministry of Education, College of Tropical Crops, Hainan University, Haikou 570228, China; Department of Plant and Microbial Biology, College of Biological Sciences, University of Minnesota, 140 Gortner Laboratory, 1479 Gortner Avenue, Saint Paul, MN 55108, USA; Systematics, Biodiversity and Evolution of Plants, Department of Biology I, Ludwig-Maximilians-Universität München, Menzinger Str. 67, 80638, Munich, Germany; Zhai Mingguo Academician Work Station, Sanya University, Sanya 572022, China; School of Integrative Plant Science, Section of Plant Biology and the L.H. Bailey Hortorium, Cornell University, Ithaca, NY 14850, USA; BTI Computational Biology Center, Boyce Thompson Institute, Ithaca, NY 14853, USA; Department of Botany, National Museum of Natural History, MRC-166, Smithsonian

**Author notes:** Authors for correspondence: Hua-Feng Wang.

**Keywords:** *Diabelia*, Phylogenomics, Gene tree discordance, Cytonuclear discordance, Hybridization

## Abstract

Understanding biological diversity and the mechanisms of the Sino-Japanese disjunctions are major challenges in eastern Asia biogeography. The Sino-Japanese flora has been broadly studied as an ideal model for plant phylogeography. *Diabelia* (Caprifoliaceae) is an East Asian genus, with a disjunctive distribution across the Sino-Japanese region. However, relationships within *Diabelia* remain elusive. In this study, we reconstructed the phylogeny of *Diabelia* and inferred historical biogeography and evolutionary patterns based on nuclear and plastid sequences from target enrichment and genome skimming approaches, respectively. We found that the main clades within *Diabelia* were discordant between nuclear and plastid trees. Both nuclear and plastid phylogenetic analyses supported five main clades: *D. serrata*, *D. tetrasepala*, *D. sanguinea*, *D. spathulata* var. *stenophylla* and *D. spathulata* var. *spathulata*. Species network analyses revealed that *Diabelia tetrasepala* is likely the result of a hybridization event. Divergence time estimation and ancestral area reconstructions showed that *Diabelia* originated in Japan during the early Miocene, with subsequent vicariance and dispersal events between Japan and Korea, and between Japan and China. Overall, our results support the division of *Diabelia* into five main clades and the recognition of five species in the genus. This research provides new insights in the species delimitation and speciation processes of taxonomically complex lineages such as *Diabelia*.

## 1 Introduction

Understanding how different factors have shaped current biological diversity is a major challenge for evolutionary biology (Jiang et al., 2016; Casebolt & Kowalewski, 2018; Wang et al., 2018; Martínez-Espinosa, 2021; Maguilla et al., 2021). The Sino-Japanese floristic region (SJFR) of East Asia was an important glacial sanctuary during the Quaternary and harbors the highest diversity of temperate plant species in the world, warranting much attention to understand its origin and diversification (Mitsui et al., 2008; Qiu et a1., 2009; 2011; Zhao et al., 2019; Tian et al. 2020; Zhang et al., 2020). The SJFR extends widely over an area from southwest China to northern Japan, with complex topography and multiple climatic zones (Qiu et a1., 2011; Lu et al., 2020). The eastern edge of the SJFR experienced dramatic changes in the palaeo-landscape during the Miocene and Pliocene (Ota, 1988; Lu et al., 2020), which may have played key roles in isolating ancient Japanese plant species from continental East Asian species.

*Diabelia* Landrein (Caprifoliaceae) is an East Asian endemic genus and was recently segregated from *Abelia* based on the paired flowers appearing at the end of short shoots (Landrein, 2010; Wang et al., 2020). This genus of shrubs belongs to the subfamily Linnaeoideae (Landrein, 2010; Wang et al., 2015; Landrein and Farjon, 2020; Wang et al., 2020) and traditionally included three species: *Diabelia serrata* (Siebold & Zucc.) Landrein with two sepals, *D. tetrasepala* (Koidz.) Landrein with four big and one smaller sepal, and *D. spathulata* (Siebold & Zucc.) Landrein with five sepals, which is further subdivided into three varieties, var. *spathulata*, var. *sanguinea* (Makino) Landrein, and var. *stenophylla* (Honda) Landrein (Fig. 1; Hara, 1983). *Diabelia* shows a disjunct distribution in the SJFR (Hara, 1983; Landrein, 2010; Zhao et al., 2019; Landrein and Farjon, 2020; Wang et al., 2020): *D. serrata* is widely distributed in southern Japan as well as on the southeastern coast of China; *D. spathulata* is distributed in south, central and northern areas of Japan, but rare in China and Korea; *D. tetrasepala* is distributed in the area from Fukushima to Fukuoka Prefectures in Japan, and its distribution range partly overlaps with that of *D. spathulata* and *D. serrata*. Based on morphological characters (the number of sepals, nectary cushion position, and corolla color), Landrein and Farjon (2020) have proposed the recognition of four species: *D. serrata*, *D. spathulata*, *D. sanguinea* (Makino) Landrein, and *D. stenophylla* (Honda) Landrein.

**Fig. 1.**
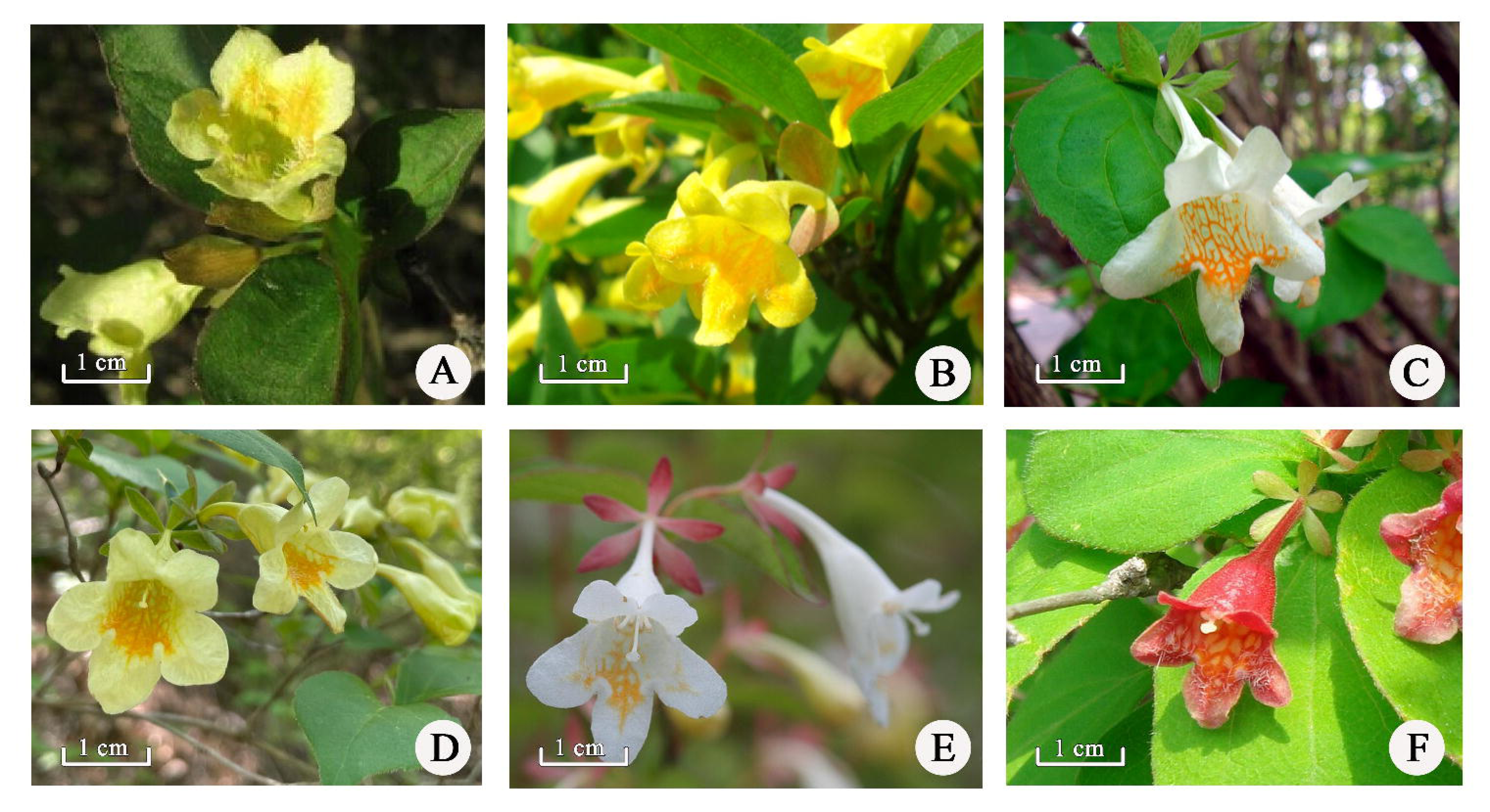
Floral diversity of *Diabelia*. (A) *Diabelia serrata* (Photograph by Difei Wu), (B) *D. serrata var. buchwaldii* (Photograph by Shota Sakaguchi), (C) *D. stenophylla* var. *tetrasepala* (Photograph by Shota Sakaguchi), (D) *D. spathulata* (Photograph by Kyoungsu Choi), (E) *D. spathulata* (Photograph by Sven Landrein), (F) *D. sanguinea* (Photograph by Sven Landrein).

There have been several previous studies on disjunct distributions between China and Japan (e.g., Lu et al., 2020; Haq et al., 2020; Takano et al., 2020), with most studies focusing on species that are widely distributed in Japan. Disjunct distribution patterns typically have been attributed to vicariance or long-distance dispersal events (Doyle et al., 2004; Baenfer et al., 2006; Bobo-Pinilla et al., 2018; Torres-Cambas et al., 2019; Wang et al., 2020; Nge et al., 2021). The disjunct distribution of *Diabelia* can be dated back to the middle Oligocene, spanning the long geological history of the formation of the flora in China and Japan (Yang et al., 2011; Shin et al., 2012; Zhao et al., 2019; Wang et al., 2020; Zhang et al., 2020). Overall, exploring the internal phylogenetic relationships and biogeographic diversification of *Diabelia* is conducive for further inference of the mechanisms leading to the disjunct distribution of SJFR in East Asia.

Several previous molecular phylogenetic studies have been conducted on *Diabelia* (Zhou et al., 2004; Landrein et al., 2012; Wang et al., 2015). However, relationships within *Diabelia* remain controversial due to limited markers and sampling. Zhou et al. (2004) conducted an Amplified Fragment Length Polymorphism (AFLP) analysis including six *Diabelia* samples and found that *D. serrata* from China was closely related to accessions of the same species from Japan. Using the nuclear internal transcribed spacer (ITS) and plastid markers (*rbcL, ndhF, matK, trnL* intron and *trnL-F* intergenic spacer), Landrein et al. (2012) found a close relationship between *D. serrata* and *D. spathulata*. Based on nine plastid markers, Wang et al. (2015) constructed a phylogeny recovering a close relationship of *D. serrata* between China and Japan. Subsequent molecular phylogenetic studies have enriched sampling across the SJFR to further investigate the phylogenetic relationships within *Diabelia*. Using plastid fragments from 37 *Diabelia* samples, Zhao et al. (2019) found the non-monophyly of several *Diabelia* species. Yet based on complete plastomes, Wang et al. (2020) recovered the monophyly of *D. serrata* while the relationships of the other species remained unresolved. Overall, sufficient informative molecular data from both nuclear and plastid genomes and a large taxonomic sampling are essential for the comprehensive reconstruction of phylogenetic relationships within *Diabelia*.

Organelle genes often exhibit phylogenetic patterns significantly different from nuclear markers (Toews and Brelsford, 2012; Leducq et al., 2017; Ji et al., 2019; Yao et al., 2019; Wang et al., 2021). Currently, target enrichment is emerging as the method of choice to obtain target sequence for numerous nuclear orthologs of many complex taxa (Wanke et al., 2017; Buys et al., 2019; Schneider et al., 2020; Granados Mendoza et al., 2020; Wang et al., 2021). Many studies have applied this method to obtain data sets for analyzing cyto-nuclear discordance, speciation, hybridization, and polyploidy (e.g., Bogarin et al., 2018; Morales-Briones et al., 2018; 2021). Given that the species-level phylogeny of *Diabelia* has remained largely unresolved, we obtained data through target enrichment and genome skimming of broadly sampled *Diabelia* accessions across the SJFR, which allowed us to (1) robustly explore intrageneric phylogenetic relationships and compare them against previous phylogenies, and (2) to elucidate the biogeography and evolution of the genus to offer a comprehensive phylogenetic framework for future studies.

## 2 Materials and Methods

### 2.1 Sampling

A data set of 47 accessions was analyzed including 42 *Diabelia* samples (encompassing all four currently recognized species of *Diabelia* and one variety of *Diabelia stenophylla* (Honda) Landrein var. *tetrasepala* (Koidz.) Landrein) (Landrein et al., 2012; Zhao et al. 2019; Landrein and Farjon, 2020; Wang et al., 2020) and five outgroup samples representing Linnaeoideae (*Abelia macrotera* var. *macrotera*, *Dipelta floribunda* Maxim., *Kolkwitzia amabilis* Graebn., *Vesalea floribunda* M.Martens & Galeotti, *Linnaea borealis* L.). Voucher specimens have been deposited in the herbarium of the Institute of Tropical Agriculture and Forestry of Hainan University (HUTB), Haikou, China. Detailed information on the geographical distribution and voucher details for the 47 samples and the localities of the populations sampled in this study are shown in Table S1.

### 2.2 DNA extraction and sequencing

We used a modified CTAB method (Doyle & Doyle, 1987) to extract total genomic DNA from silica gel-dried tissue or herbarium accessions. The concentration of each extraction was checked with a Qubit 2.0 Fluorometer (Thermo Fisher Scientific, Waltham, MA, USA). A total of 400 ng of DNA was sonicated with a Covaris S2 (Covaris, Woburn, MA) to produce fragments ∼150-350 bp in length prior to library preparation. Libraries of genomic DNA were made following Weitemier et al. (2014). To ensure that genomic DNA was sheared to the appropriate fragment size, all samples were evaluated on a 1.2% (w/v) agarose gel.

To obtain nuclear data we used a target enrichment approach (Weitemier et al., 2014) and plastid data we used genome skimming (Wang et al., 2020). Baits designed across Dipsacales (Wang et al., 2021) were used to target 428 putatively single-copy genes. Hybridization, enrichment, and sequencing followed Wang et al. (2021).

### 2.3 Reads processing and assembly

Trimmomatic v.0.36 (Bolger et al., 2014) was used to remove adaptor sequences and low-quality bases from the raw reads (ILLUMINACLIP: TruSeq_ADAPTER: 2:30:10 SLIDINGWINDOW: 4:5 LEADING: 5 TRAILING: 5 MINLEN: 25). HybPiper v.1.3.1 (Johnson et al., 2016) was employed to assemble the nuclear loci. Exons were assembled individually to avoid chimeric sequences in multi-exon genes produced by potential paralogy (Morales-Briones et al., 2018), in which exons longer than 150 bp were set as a reference. Paralog detection was undertaken for all exons using the ‘paralog_investigator’ option in HybPiper. To obtain ‘monophyletic outgroup’(MO) orthologs (Yang and Smith, 2014), all assembled loci (with and without paralogs detected) were processed following Morales-Briones et al. (2021).

For the plastome assemblies, clean reads were extracted from the raw sequencing reads by using SOAPfilter_v2.2 to remove adapter sequences and low-quality reads. The resulting reads were used as the input to assemble plastomes using MITObim v1.8 (Hahn et al. 2013) following Wang et al. (2020).

### 2.4 Phylogenetic analyses

#### Nuclear data sets

We used concatenation and coalescent-based methods to analyze the nuclear data. For concatenation analyses, sequences of individual nuclear exons were aligned with MAFFT v.7.407 (Katoh & Standley, 2013) and columns with more than 90% missing data were removed using Phyutility (Smith and Dunn, 2008). A maximum likelihood tree from the concatenated matrix was inferred with RAxML v.8.2.20 (Stamatakis, 2014) using a partition-by-locus scheme and the GTRGAMMA substitution model for all partitions. We assessed clade support with 100 rapid bootstrap replicates (BS). For coalescent species tree estimation, ASTRAL-III v.5.7.1 (Zhang et al., 2018) was used to estimate a species tree based on individual exon trees constructed using RAxML with a GTRGAMMA model. We used local posterior probabilities (LPP) to calculate branch support (Sayyari and Mirarab, 2016). To evaluate nuclear gene tree discordance, we used Quartet Sampling (QS; Pease et al., 2018) to distinguish strong conflict from weakly supported branches with 1000 replicates. Additionally, we calculated the internode certainty all (ICA) score (Salichos et al., 2014) and the number of conflicting and concordant bipartitions on each node of the species trees with Phyparts (Smith et al., 2015).

#### Plastome data set

Complete plastomes were aligned using MAFFT v.7.407 (Katoh and Standley, 2013). A ML tree was inferred with IQ-TREE v.1.6.1 (Nguyen et al., 2015) under the extended model and 200 non-parametric BS replicates for branch support. In addition, branch support was evaluated using QS with 1000 replicates.

### 2.5 Species network analysis

We used PhyloNet (Wen et al., 2018) to infer a maximum pseudo-likelihood species network with the InferNetwork_MPL command (Yu and Nakhleh, 2015). Due to computational restrictions and given that we were primarily concerned with the underlying network between different species, we reduced the 47-taxon data set to one outgroup and 14 ingroup taxa to represent all major clades within *Diabelia*. Network searches were undertaken using only nodes in the gene trees that had at least 50% bootstrap support, allowing up to five hybridization events while optimizing branch lengths and inheritance probabilities of the returned species networks under the full likelihood. The command ‘CalGTProb’ in PhyloNet was used to infer the maximum likelihood of the concatenated RAxML, ASTRAL, and plastid trees, given the individual gene trees to estimate the optimal number of hybridization events and inspect whether the species network represented a better model than a purely bifurcating tree. The bias-corrected Akaike information criterion (AICc; Sugiura, 1978), Akaike information criterion (AIC; Akaike, 1973), and Bayesian information criterion (BIC; Schwarz, 1978) were used for model selection, with the best-fit-species network having the lowest AICc, AIC and BIC scores.

### 2.6 Divergence time estimation

Divergence times were estimated with BEAST v.2.4.0 (Bouckaert et al., 2014) using the concatenated nuclear data set. The root age was set to 50 Ma (lognormal prior distribution 32.66 - 57.81 Ma) following Wang et al. (2021). We selected the fruit fossil of *Diplodipelta* Manchester & Donoghue (Bell & Donoghue, 2005) that could confidently be placed in our tree as an internal calibration point, with the constraint set to 36 Ma (offset 36 Ma, lognormal prior distribution 34.07 - 37.20 Ma). Dating analyses were carried out using an uncorrelated lognormal relaxed clock under the GTR + G model for each marker partitioned separately and a Yule tree prior. The MCMC chains were run for 500,000,000 generations and sampling every 5,000 generations. Tracer v.1.7 (Drummond et al., 2012) was used to check convergence with the first 10% of trees removed as burn-in and to assess that all effective sample size (ESS) values were ≥ 200. The produced tree files were combined using LogCombiner v1.8.2 and the maximum clade credibility tree was generated in TreeAnnotator v1.8.4 (Drummond et al., 2012).

### 2.7 Ancestral area reconstruction

The ancestral area reconstruction was done using the Statistical Dispersal- Vicariance Analysis (S-DIVA) in RASP version 4.2 (Yu et al., 2015) based on Bayesian Binary Method (BBM) with the concatenated nuclear data set. To prevent biased inferences towards wide or unlikely distributions for the crown node of the ingroup (only *Diabelia* species) due to the uncertainty in the root area of the outgroup, we pruned the five outgroups for our ancestral state reconstructions. Three areas were defined to cover the present distribution range of *Diabelia*: (A) Japan, (B) Korea, and (C) China. Distribution areas of all populations in this study were defined according to field observations (Table S1). The BBM analyses ran for 500,000,000 generations using 10 MCMC chains.

### 2.8 Data accessibility

Raw Illumina data from target capture are available in the Sequence Read Archive (SRA) under accession SUB10211626 (see Table S1 for individual sample SRA accession numbers). DNA alignments, phylogenetic trees and results from all analyses and data sets can be found in the Dryad data repository XXXXXX.

## 3 Results

### 3.1 Exon assembly

The number of assembled exons per species (with > 75% of the target length; except the outgroups) ranged from 587 (*D. serrata* E314) to 989 (*D. sanguinea* E301), with an average of 889 exons per sample (Table S2; Fig. S1). The number of exons with paralog warnings (except the outgroups) ranged from 34 in *D. serrata* E314 to 619 in *D. sanguinea* E301 (Table S2). The concatenated alignment had a length of 392,600 bp, including 18,057 parsimony-informative sites, and a matrix occupancy of 86.7% (Table 1). The plastid alignment resulted in a matrix of 159,365 bp with 4,693 parsimony-informative sites and a matrix occupancy of 98.2% (Table 1).

**Table 1.**
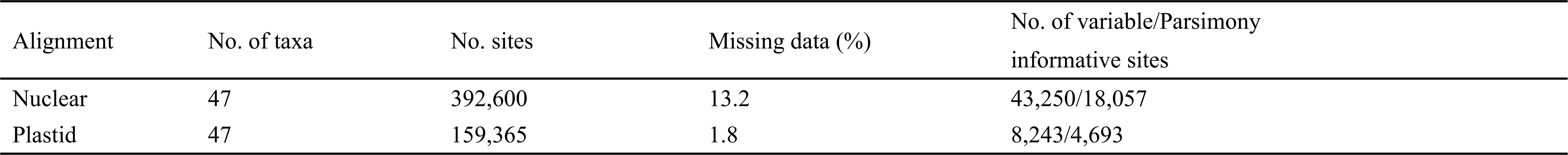
Data set statistics, including the number of taxa, number of characters, number of PI characters, missing data.

### 3.2 Phylogenetic reconstruction

Our nuclear phylogenetic analyses recovered the monophyly of *Diabelia* with maximum support (BS = 100; LPP = 1) and recognized five main clades: *D. serrata*, *D. stenophylla* var. *tetrasepala*, *D. sanguinea*, *D. stenophylla* and *D. spathulata*. However, the relationships of the five main groups within *Diabelia* varied among analyses and data sets.

With respect to the nuclear concatenated ML tree, the phylogeny was recovered with moderate (50 < BS ≤ 70) to high (BS > 70) support along the five main clades (Figs. 2, S3 - S4). The monophyly of *Diabelia* was supported by 313 gene trees (out of 439 informative gene trees; ICA = 0.41) and full QS support (1.0/–/1.0; i.e., all sampled quartets supported that node). *Diabelia serrata* forms a clade with strong QS support (0.91/0.4/1) and only 34 concordant trees (out of 427; ICA = 0.02). The monophyly of *D. stenophylla* var. *tetrasepala* was supported by 158 out of 316 informative trees (ICA = 0.24) and full QS support. The sister relationship of *D. stenophylla* var. *tetrasepala* and *D. serrata* was supported by only 12 gene trees (out of 478; ICA = 0.01) and had moderate QS support with a signal of a possible alternative topology (0.21/0.35/0.99). *Diabelia sanguinea* was supported by only 24 gene trees (out of 395; ICA = 0.03) but had full QS support. The sister relationship of *D. sanguinea* and the clade of *D. serrata* + *D. stenophylla* var. *tetrasepala* was supported by only 4 gene trees (out of 490; ICA = -0.02) and moderate QS support, with a signal for a possible alternative topology (0.21/0.49/0.99). The clade of *D. stenophylla* had 43 concordant trees (out of 376; ICA =0.04) and high QS support (0.91/0.33/1). *Diabelia spathulata* was supported by 26 gene trees (out of 304; ICA = 0.06) and moderate QS support (0.22/0.29/0.98), with the sister relationship to the remainder of *Diabelia*.

**Fig. 2.**
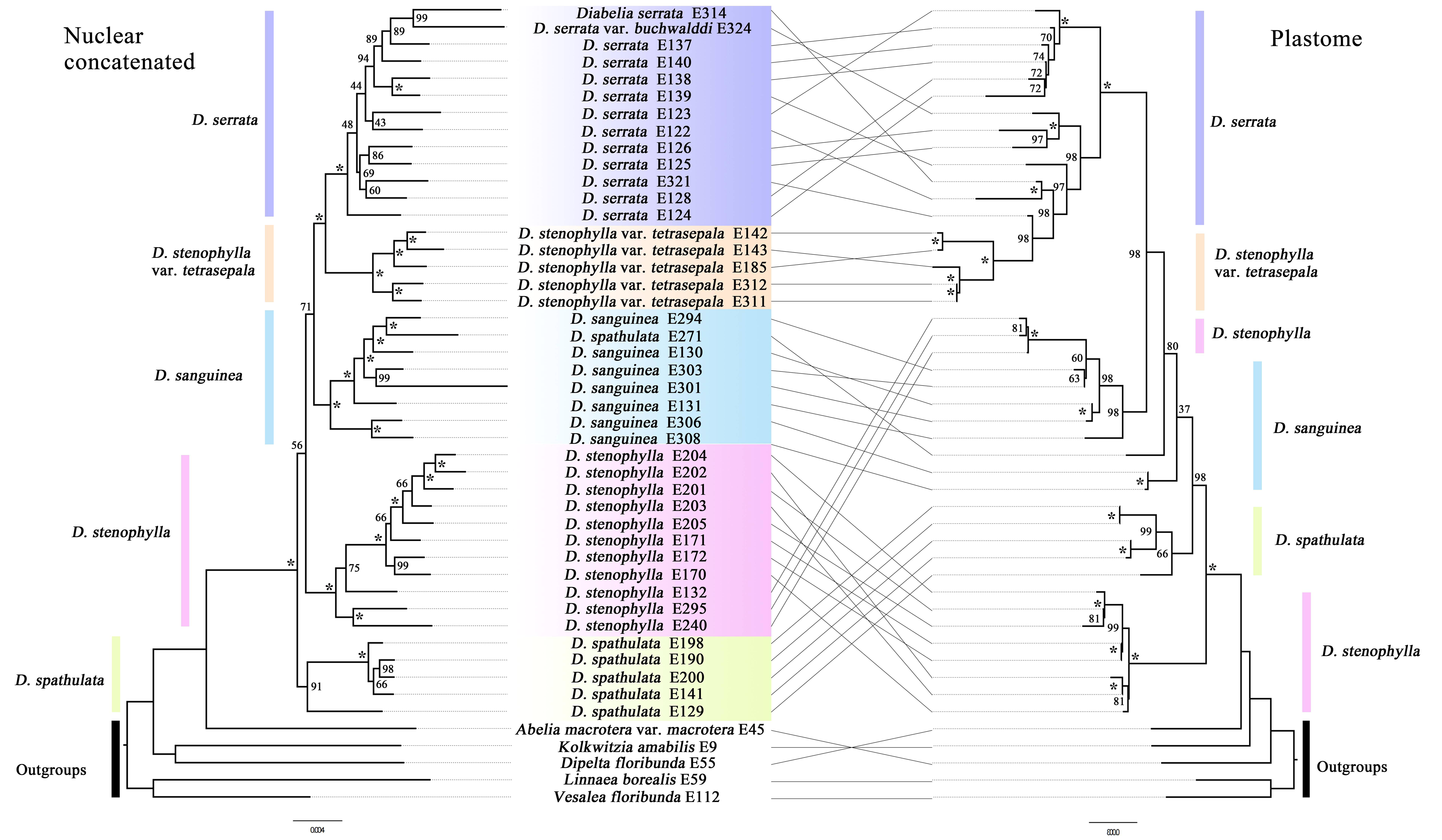
Tanglegram of the nuclear concatenated (left) and plastid (right) phylogenies of *Diabelia*. Dotted lines connect taxa between the two phylogenies. Maximum likelihood bootstrap support values are shown above branches. The asterisks indicate maximum likelihood bootstrap support of 100%. Major taxonomic groups or main clades in the family as currently recognized are indicated by branch colors as a visual reference to relationships.

The ASTRAL species tree (Fig. 3, S2) presented a largely congruent topology compared to the nuclear ML tree with all five major groupings recovered, each being respectively monophyletic. The main clades were highly supported (LPP ≥ 0.93) except for the sister node of *D. sanguinea* and *D. serrata* (LPP = 0.44). The monophyly of *D. serrata* had strong QS support with a signal for a possible alternative topology (0.92/0.38/1). *Diabelia spathulata* var. *sanguinea* was supported by moderate QS support (0.62/0.78/0.99). The sister relationship of the clade of *D. sanguinea* and *D. serrata* was supported by only four gene trees (out of 496; ICA = - 0.02) and had counter QS support with signal for a possible alternative topology (-0.22/0.21/0.99). *Diabelia stenophylla* var. *tetrasepala* was sister to the clade of *D. serrata + D. sanguinea* with moderate QS support (0.2/0.35/0.99). The clade of *D. stenophylla* between China and Japan had strong QS support (0.9/0.2/1).

**Fig. 3.**
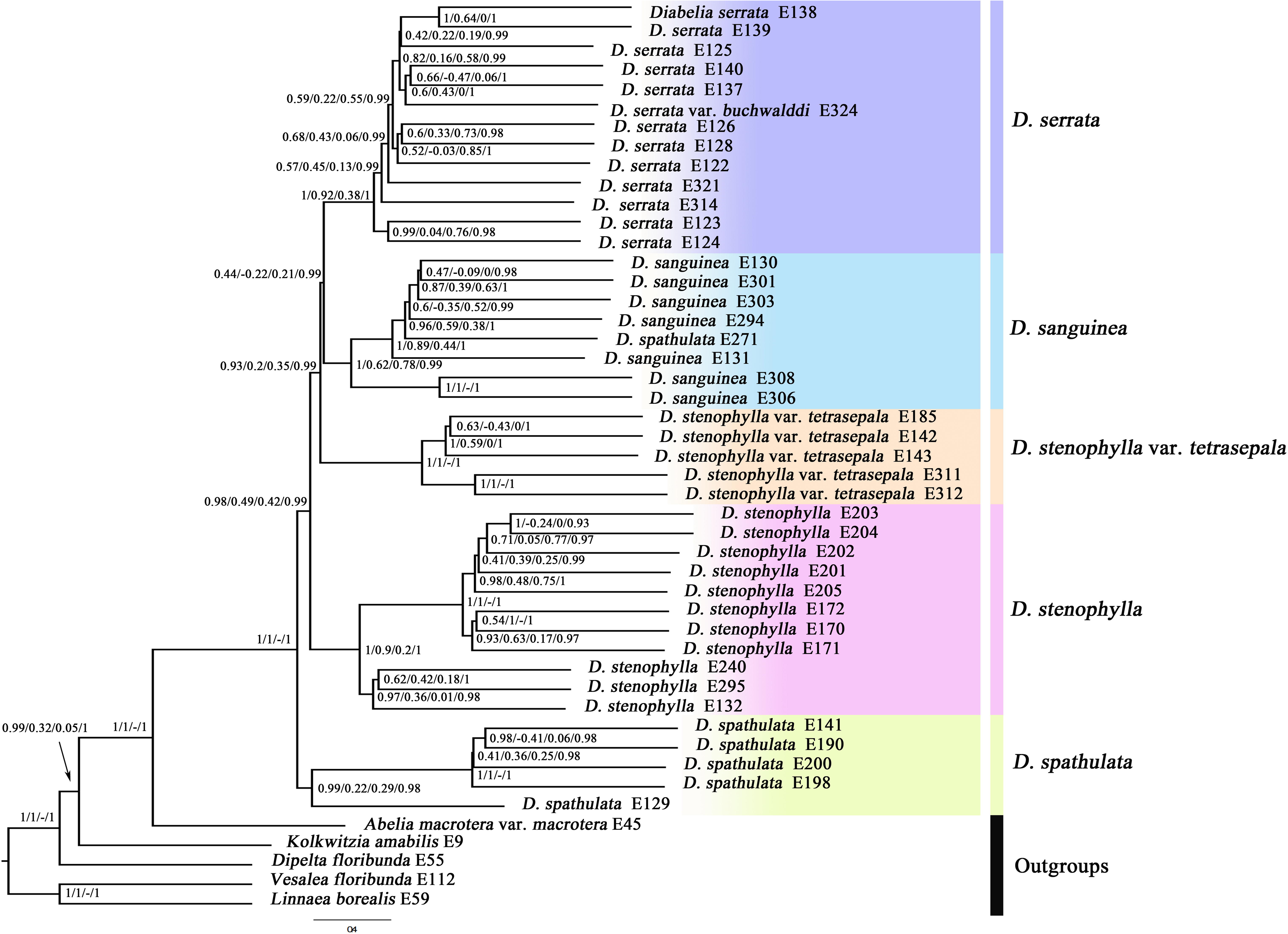
ASTRAL species tree of *Diabelia*; Clade support is depicted as: Local posterior probability (LPP)/Quartet concordance (QC)/Quartet differential (QD)/Quartet informativeness (QI).

The phylogenetic relationships in the plastid ML tree of *Diabelia* (Fig. 2 and S5) were also confirmed with full QS support. The clade of *D. sanguinea + D. stenophylla* (Japan) was sister to the *D. serrata + D. stenophylla* var. *tetrasepala* clade, and the combined clade had strong support (BS = 98) and moderate QS score (0.14/0.57/0.98). There were some observed topological differences between the plastid and nuclear trees. For example, the plastid trees showed that *D. stenophylla* var. *tetrasepala* was nested within *D. serrata* with strong support (BS = 98) and full QS support; *D. sanguinea* was not-monophyletic; and *D. stenophylla* from Zhejiang was *s*ister to the remainder of *Diabelia*, rather than to *D. spathulata*.

### 3.3 Species network analysis

The species network analyses (15-taxa data set) recovered topologies with one to five reticulation events, which appear to be a better model than a strictly bifurcating tree (Table 2; Fig. S6). Model selection preferred the network with two reticulation events (Fig. 4). With this preferred network, the first reticulation event involved *D. serrata* and *D. spathulata*. The inheritance probabilities of this event showed that the ancestral lineage of *D. stenophylla* var. *tetrasepala* had the largest genomic contribution of 65.9 % from the clade of *D. spathulata*, and a smaller portion (34.1 %) came from the *D. serrata* clade. Another reticulation event was observed indicating the clade of *D. spathulata* + *D. stenophylla* had genetic contributions (79.3%) from the lineage leading to *D. sanguinea*.

**Fig. 4.**
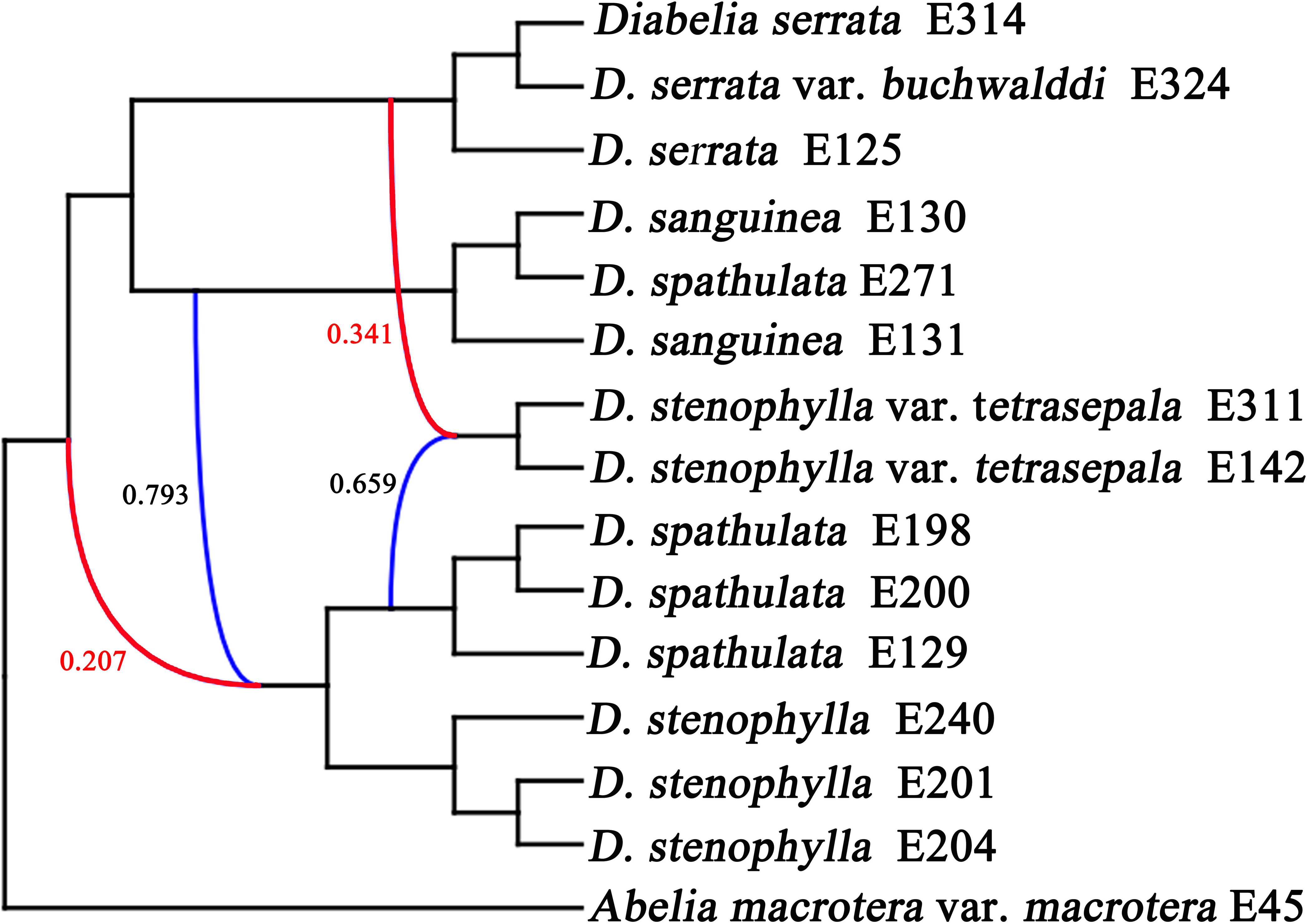
Best supported species network inferred with PhyloNet for the 15-taxa. Numbers next to the inferred hybrid branches indicate inheritance probabilities. Blue curved lines represent major hybrid edges. Red curved lines represent minor hybrid edges (edges with an inheritance contribution < add space 0.50).

**Table 2.**
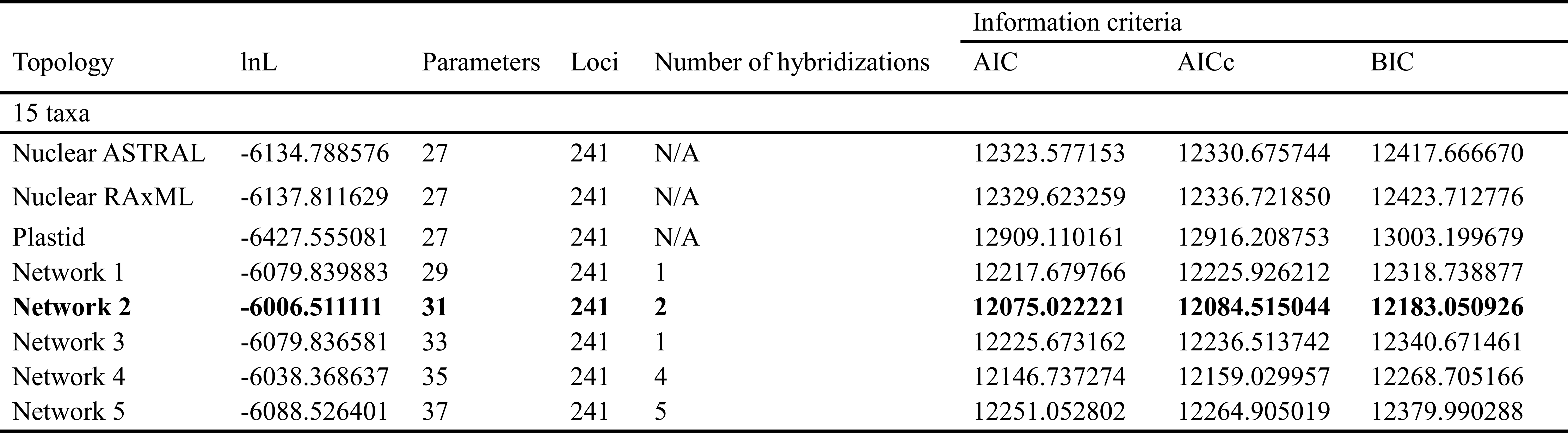
Model selection of the different species networks and bifurcating trees.

### 3.4 Divergence time estimation

Our dating estimates based on the nuclear data suggested an early Miocene crown age for *Diabelia* of 22.53 Ma (95% Highest Posterior Density (HPD): 17.89 - 27.73 Ma, node 1 in Fig. 5). The sister relationships of *D. serrata* and *D. stenophylla* var. *tetrasepala* diverged during the early Miocene 18.17 Ma (95% HPD: 14.07 - 22.41 Ma, node 4 in Fig. 5). The crown age of *D. serrata* was dated to 14.88 Ma (95% HPD: 11.10 - 18.91 Ma, node 5 in Fig. 5). The split between *D. spathulata* (E271) and *D. sanguinea* occurred between 3.90 and 9.41 Ma (node 7 in Fig. 5). The diversification of *D. spathulata* between Korea and Japan was estimated as 4.78 Ma (95% HPD: 3.02 - 6.84 Ma, node 8 in Fig. 5). The divergence time of *D. stenophylla* between China and Japan was 6.30 Ma (95% HPD: 3.90 - 9.41 Ma, node 6 in Fig. 5).

**Fig. 5.**
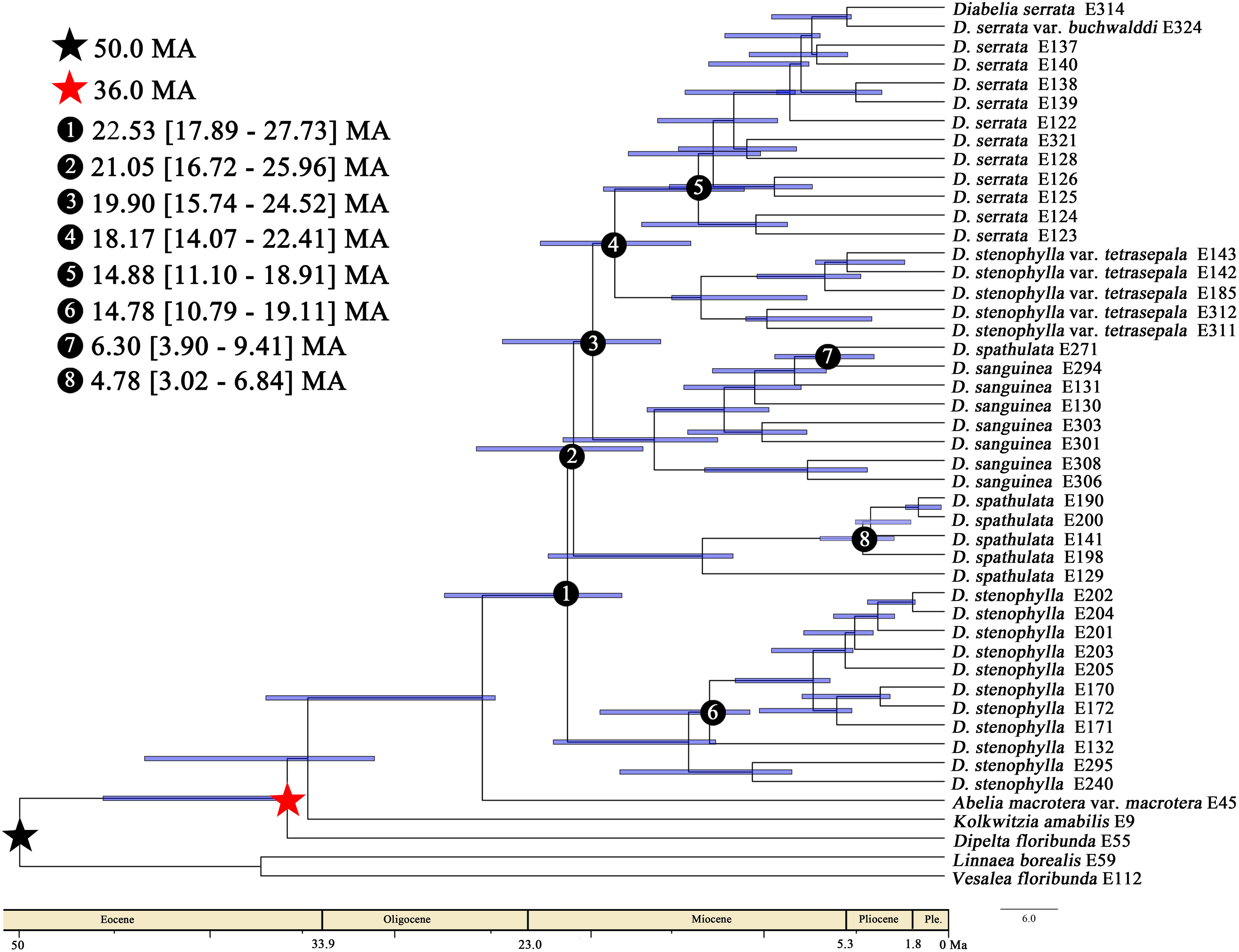
BEAST analysis of divergence times based on the nuclear data set. Calibration points are indicated by A, B. Numbers 1-8 represent major divergence events in *Diabelia*; mean divergence times and 95% highest posterior densities (HDP) are provided for each node of interests. Blue bars represent 95% HDP.

### 3.5 Ancestral area reconstruction

The reconstruction results of the ancestral distribution of *Diabelia* were presented in Fig. 6. There were three dispersal events and three vicariance events identified within the three defined biogeographic areas (Fig. 6). Our analyses revealed that the ancestor of *Diabelia* was present throughout Japan (region A) in the early Miocene (Fig. 6), followed by dispersal or vicariance events across Japan, Korea, and China. Migration events occurred primarily during the Neogene. *Diabelia spathulata* from Korea was suggested to have originated in Northeast Japan. However, one vicariance and one dispersal event were detected for the ancestral nodes of *D. stenophylla* from China, supporting their origin from Japan.

**Fig. 6.**
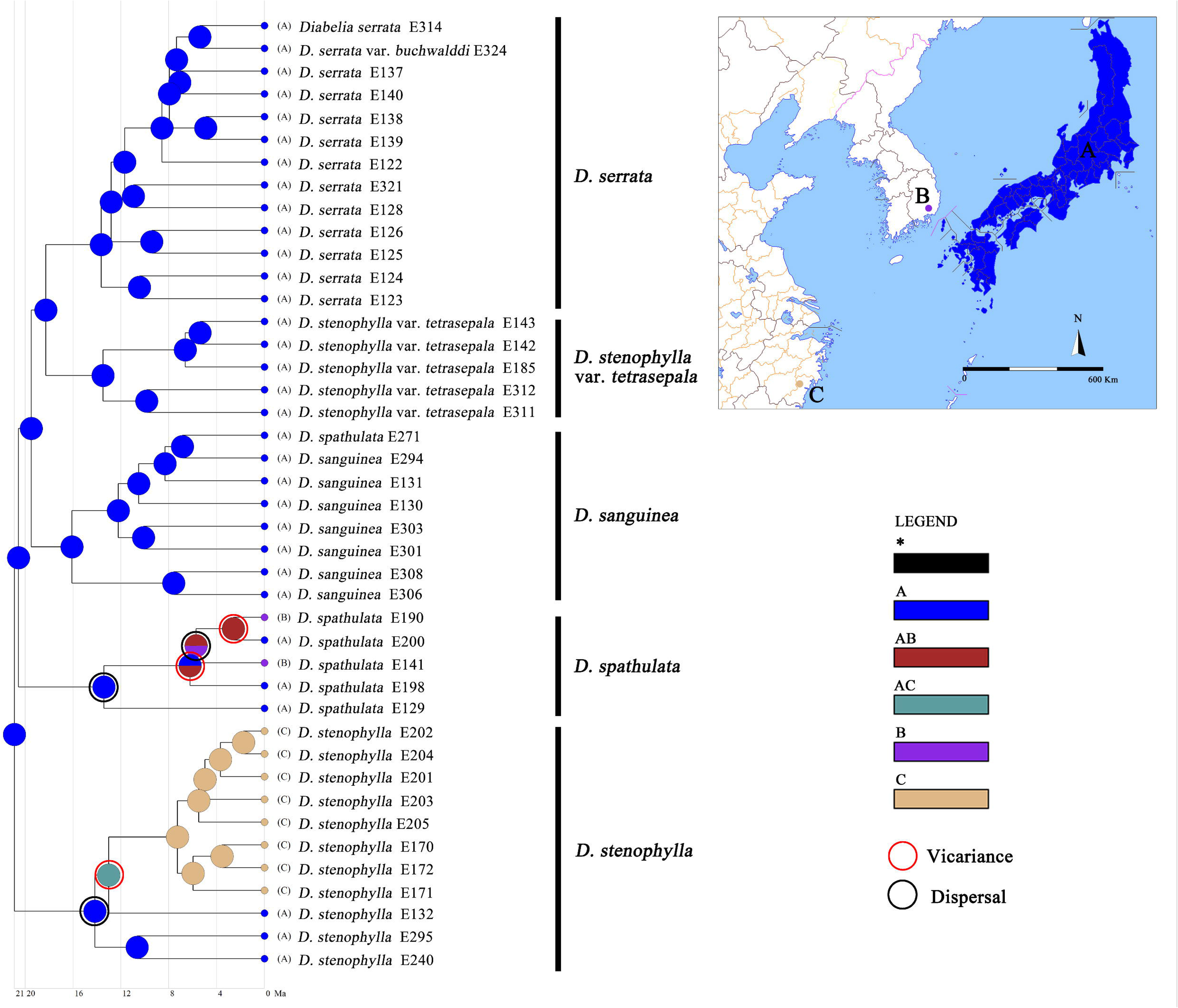
Ancestral area reconstruction for *Diabelia*. Areas of endemism are as follows: (A) Japan, (B) Korea, (C) China. The numbered nodes represent crown nodes of important colonization events.

## 4 Discussion

### 4.1 Phylogenetic incongruence and hybridization, and implications for species delimitation

Our phylogenetic analyses retrieved the same five main clades of *Diabelia* in the nuclear ML tree, which are also supported in the ASTRAL tree, but the relationships among these clades are incongruent between the nuclear and plastid data (Figs. 2 and 3). For example, *D. serrata* and *D. stenophylla* var. *tetrasepala* were recovered as monophyletic in the nuclear ML tree, while *D. stenophylla* var. *tetrasepala* was nested within *D. serrata* in the plastid ML tree (Fig. 2). We not only detected widespread cytonuclear discordance across *Diabelia* (Fig. 2), but our results also showed extensive conflict among individual gene trees and the species trees (Figs. 3 and S2). Several processes can lead to gene tree inconsistencies between closely related groups, including incomplete lineage sorting (ILS), hybridization, horizontal gene transfer, or gene duplication and loss (Morales-Briones et al., 2018; Bogarin et al., 2018; Carter et al., 2019; Stull et al., 2020; Wang et al., 2021). Ancestral polymorphisms may lead to incomplete genealogical classification, therefore phylogenetic relationships between organelle markers may fail to capture the true process of population differentiation, or inconsistencies in gene trees may reflect interspecies hybridization and cytoplasmic infiltration (Lee-Yaw et al., 2019; Wielstra & Arntzen, 2020; Tkach et al., 2020). Therefore, exploring the causes of inconsistencies in gene trees may help explain the relative influences of drift, gene flow and selection on the maintenance of organelle variation within and between groups, which is helpful for revealing the evolutionary process of inter-group relationships (Mao et al., 2020; Dufresnes et al., 2020).

Comparing the topologies between the nuclear and plastid data, we found that the topology from the nuclear gene data was more stable. The nuclear data recovered five strongly supported clades (Figs. 2 and 3), with *D. serrata* forming a stable monophyletic group that was consistent with previous studies supporting the monophyletic nature of *D. serrata* (Wang et al., 2020). *Diabelia stenophylla* and *D. spathulata* were each also recovered as monophyletic based on the nuclear data. While *D. sanguinea* was not recovered as a monophyletic group in the plastid tree (Fig. 2), the possibility exists that the non-monophyly of *D. sanguinea* is due to plastid capture events or ILS (Liu et al., 2017; Renoult et al., 2009; Wang et al., 2020). Notably, the placement of *D. sanguinea* in the nuclear data had full LPP support with low ICA value and moderate QS score, which suggests that ILS and/or unidentified hybrid lineages continue to obscure our understanding of the relationship of *D. sanguinea* within the genus, and even the relationships of *Diabelia* as a whole, due to strong signals of gene tree discordance. Previous phylogenetic relationships of *Diabelia* based on only plastid data have also been unstable (Zhao et al., 2019; Wang et al., 2020). In the present study, the nuclear data provide a more robust estimate of the species tree compared to plastid data, suggesting that sufficiently informative molecular data are important to our understanding of relationships in *Diabelia*.

A previous study (Zhao et al., 2019) reported gene introgression events between *Diabelia* species and speculated that *D. stenophylla* var. *tetrasepala* (four big and one small sepals) may have resulted from hybridization between *D. serrata* (two sepals) and *D. spathulata* (five sepals). However, this hypothesis has not been confirmed. In this study, we conducted network analyses for *Diabelia* and our results support the existence of reticulation events within the genus (Fig. 4). Concerning the reticulate evolution of the *D. stenophylla* var. *tetrasepala* clade, the inheritance contributions (34.1 % and 65.9 %) support the hybridization event between *D. serrata* and *D. spathulata.* Based on two similar morphologies but different origins of taxa within *Melastoma*, Zou et al. (2017) suggested that it is difficult to infer the origins of hybrid taxa based only on morphology, and the hybrids may be the result of small range overlaps among parental species. Morales-Briones et al. (2018), based on the extensive history of hybridization and network results in *Lachemilla*, showed the potential of phylogenetic species network methods to investigate phylogenetic discordance caused by ILS and hybridization. Given our network results and the distribution of *D. stenophylla* var. *tetrasepala* partly overlaps with those of parental populations, we argue that *D. stenophylla* var. *tetrasepala* may be of hybrid origin between *D. serrata* and *D. spathulata*. As noted by Morales-Briones et al. (2018), additional biological information is necessary for a robust interpretation of values from the phylogenetic networks. Hence, we will include more samples to obtain a more robust inference of relationships for future studies.

Despite the strong signals of gene tree discordance, our nuclear and plastid phylogenies strongly supported five major clades in *Diabelia*: *Diabelia serrata*, *D. stenophylla* var. *tetrasepala*, *D. spathulata*, *D. sanguinea*, and *D. stenophylla* (Figs. 2, 3). Especially in the nuclear data, all five major branches are monophyletic. These correspond to the species of *D. serrata*, *D. spathulata*, *D. sanguinea*, and *D. stenophylla*, but the clade of *D. stenophylla* var. *tetrasepala* does not form a monophyletic lineage with *D. stenophylla*. To resolve the non-monophyly of *D. stenophylla*, and to clarify the phylogenetic relationship within *Diabelia*, we raise *D. stenophylla* var. *tetrasepala* to the species level, recognizing it as *Diabelia tetrasepala* (Hara, 1983; Landrein, 2010). Hara (1983) also recognized *Diabelia tetrasepala* as a distinct species. Additionally, this taxon can be easily distinguished by the form of sepals (sepals 4 plus a reduced adaxial sepal lobe in *D. tetrasepala*, sepals 5 of similar size in *D. stenophylla*) (Zhao et al., 2019; Landrein and Farjon, 2020; Wang et al., 2020). Overall, based on our phylogenetic inferences (nuclear), we recognize five species in the genus (i.e., *D. serrata*, *D. tetrasepala*, *D. spathulata*, *D. sanguinea*, and *D. stenophylla*).

### 4.2 Molecular dating and demographic analyses

Based on our nuclear chronogram (Fig. 5), all major *Diabelia* lineages differentiated 22.53-18.17 Ma (Fig. 5). Notably, this point estimate broadly coincides with the isolation of the Japanese island (24-22 Ma) (Hotta, 1974; Iijima and Tada, 1990; Li et al., 1996; Maekawa, 1998) and the estimated dates are earlier than previous estimates based on chloroplast data (Wang et al., 2020). The East China Sea (ESC) land bridge likely acted as a barrier to the dispersal of plant species during the LGM and earlier cold periods, despite its repeated exposure during the Miocene (7.0-5.0 Ma) and Quaternary (2.0-1.3 Ma, 0.2-0.015 Ma) (Harrison, 2001). Even though the Japanese archipelago was not covered by a major ice sheet during the last glacial period (Ono, 1984), the mean annual temperature was 5℃-9℃ cooler and the precipitation was less than at present (Yasuda and Narita, 1981; Tsukada, 1988). In addition, because of lower sea levels (ca. 100 m below present), Shikoku and Kyushu were continuous with Honshu, and the continental shelf, ca. 20-30 km from the present coastline, emerged around the archipelago (Ohta and Yonekura, 1987). The distribution data of *D. serrata* suggests a wide distribution in Japan except for Hokkaido. As the climate warmed, the species recovered and expanded northward or towards higher altitudes (Tsukada, 1988; Takahara et al., 2000), therefore, forming the current isolated distribution.

*Diabelia* species are relatively tolerant to cold and arid climates, which might have facilitated gene exchange across the glacially exposed ECS land bridge up until its latest submergence. All *Diabelia* species have fruits forming samaras, which increase long-distance dispersal abilities (Landrein, 2010). Our molecular dating indicates that *Diabelia* originated in the Japanese archipelago (Fig. 5) and the onset of diversification of *D. stenophylla* from Japan and *D. stenophylla* from China occurred during the early Pliocene. Our results suggest a vicariance event and a dispersal event associated with *D. stenophylla* from China (Fig. 6). Under the hypothesis of long-distance dispersal, a predicted lineage in one region should nest within a lineage from a separate disjunctive region. In contrast, under vicariance, lineages from different geographic regions would each be monophyletic with relatively comparable levels of genetic diversity in each region of the distribution (Yoder & Nowak, 2006; Liao et al., 2016; Thomas et al., 2017). The disjunction between *D. stenophylla* in Japan and *D. stenophylla* from China may have resulted from migration across this land bridge followed by vicariance.

Our results show that that the interchanges of the populations of *D. spathulata* between Korea and Japan are frequent. The close relationships between Japan and Korea have also been observed in other taxa such as *Meehania urticifolia* (Takano et al., 2020) and *Kirengeshoma koreana* (Qiu et a1., 2011). Furthermore, our results also suggest that populations of *D. spathulata* in Korea originated from Japan and are likely due to recent vicariance and dispersal events (Fig. 6), which is largely congruent with the previous conclusions by Wang et al. (2020). Wang et al. (2020) showed that the populations between Korea and the northern Japan may have resulted from a vicariance event. Our results further suggest an early and a later dispersal event, supporting a highly dynamic biogeographic relationships between Japan and Korea. Our divergence time estimates suggest that species differentiation in *D. spathulata* occurred during the late Miocene to the early Pliocene (4.78 Ma, 95% HPD: 3.02–6.84 Ma), suggesting that extant populations likely differentiated well before the LGM. The cooling and drying climate in the late Miocene drove the formation of *D. spathulata*. The changing climate in the late Pliocene and Pleistocene is related to lineage differentiation, genetic diversity, and population contraction and expansion, which have also been observed in *Cercidiphyllum japonicum* (Qi et al., 2012) and *Euptelea* (Cao et al., 2016). This study adds another example of this well-documented pattern.

## 5 Conclusions

A robust phylogeny of *Diabelia* was reconstructed with nuclear and plastid data based on target enrichment and genome skimming approaches. The inferred phylogenies from both nuclear and plastome data indicate that *Diabelia* can be divided into five main clades, which are further supported by morphological traits such as number of sepals, nectary cushion position, and corolla color. Our results show clear cytonuclear discordance and strong conflict between individual gene trees and species trees in *Diabelia*. The PhyloNet results further confirmed the existence of reticulation events in *Diabelia*, supporting that *D. tetrasepala* was the result of a hybridization event. The divergence time and biogeographic analyses further support the differentiation and propagation of *Diabelia* with multiple vicariance events from the perspective of time and space, further supporting the complex natural hybridization and evolutionary network of the disjunctive flora of Japan and China. Tree-like and reticulate evolution should be considered when reconstructing phylogenetic relationships among closely related species. The appropriate choice of data to construct phylogenetic trees is important in the era of genomics. Further studies are needed to clarify the origin, dispersal and evolution of Sino-Japanese disjunct species with nuclear and plastid data at the population level. Finally, our results shed light on the species delimitation, supporting the recognition of five species in *Diabelia*, corresponding to the five main clades within the genus in the nuclear phylogeny.

### Taxonomic treatment

Based on our results, we formally recognize five species in *Diabelia*. Landrein & Farjon (2019) treated four of the five species, i.e., *D. sanguinea*, *D. serrata*, *D. stenophylla*, and *D. spathulata*. Below we provide the synoptic treatment for *Diabelia tetrasepala*.

***Diabelia tetrasepala*** (Koidz.) Landrein, Phytotaxa 3: 37. 2010.

Basionym: *Abelia spathulata* Siebold & Zucc. var. *tetrasepala* Koidz., Bot. Mag. (Tokyo) 29 (348): 311. 1915.

*Abelia tetrasepala* (Koidz.) H. Hara & S. Kuros. in Kurosawa & Hara, J. Jap. Bot. 30: 296. 1955; *Diabelia stenophylla* (Honda) Landrein var. *tetrasepala* (Koidz.) Landrein, Kew Bull. 74(4)-70: 186. 2019.

*Abelia spathulata* Siebold & Zucc. var. *subtetrasepala* Makino, J. Jap. Bot. 1: 18. 1917.

*Abelia spathulata* Siebold & Zucc. var. *subtetrasepala* Makino f. *flavescens* Honda, J. Jap. Bot. 11: 569. 1935.

## Acknowledgements

The work was funded by Hainan Provincial Natural Science Foundation of China (421RC486) and National Natural Science Foundation of China (31660055). We thank Gabriel Johnson for his help with the target enrichment experiment, and the United States National Herbarium for permission to sample some collections. We acknowledge the staff in the Laboratories of Analytical Biology at the National Museum of Natural History, the Smithsonian Institution for support and assistance.

## Author contributions

H.-F.W. and J.W. conceived the study. H.-F.W. and D.F.M-B. performed the research and analyzed the data. X.-R.K., H.-F.W., D.F.M-B., J.W., H.-X.W., Q.-H.S. and J.B.L. wrote and revised the manuscript.

## Figure Legends

**Fig. S1.**
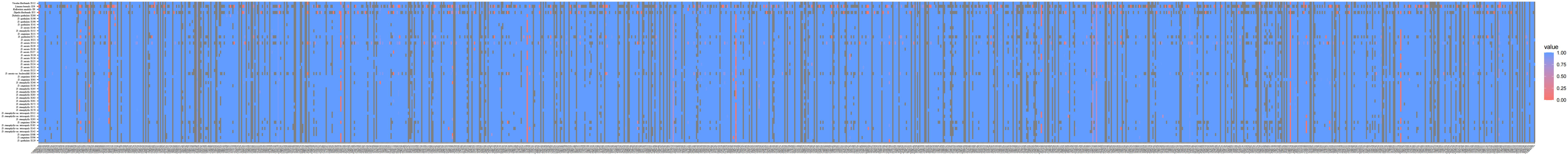
Heatmaps showing gene recovery efficiency for the nuclear genes in 42 species of *Diabelia*. Columns represent genes, and each row is one sample. Shading indicates the percentage of the reference locus length coverage.

**Fig. S2.**
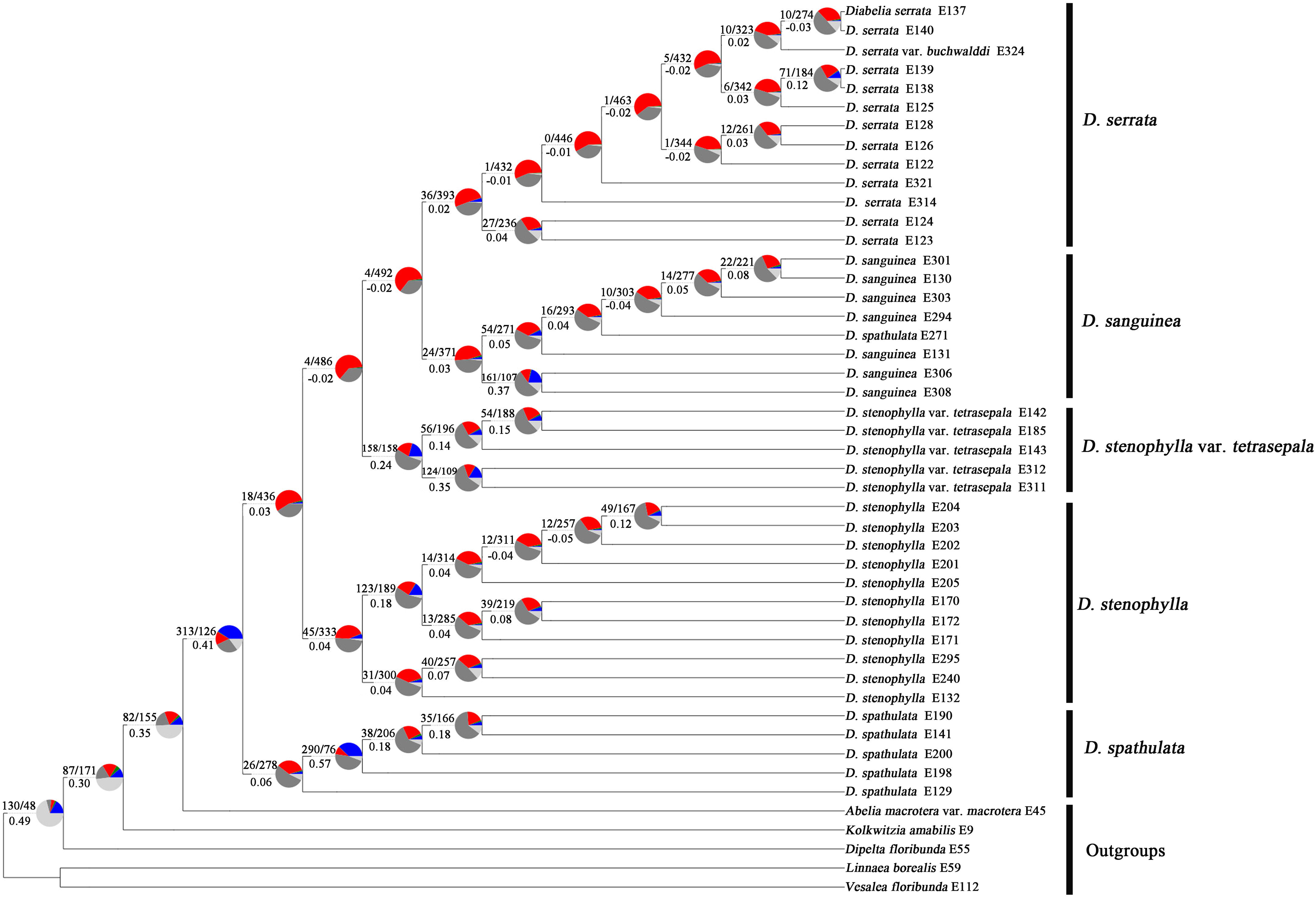
ASTRAL species tree. Numbers above branches indicate the number of gene trees concordant/conflicting with that node in the species tree. Numbers below the branches are the Internode Certainty All score. Pie charts next to the nodes present the proportion of gene trees that supports that clade (blue), the proportion that supports the main alternative for that clade (green), the proportion that supports the remaining alternatives (red), light gray means missing data, and dark gray means uninformative (BS < 50%).

**Fig. S3.**
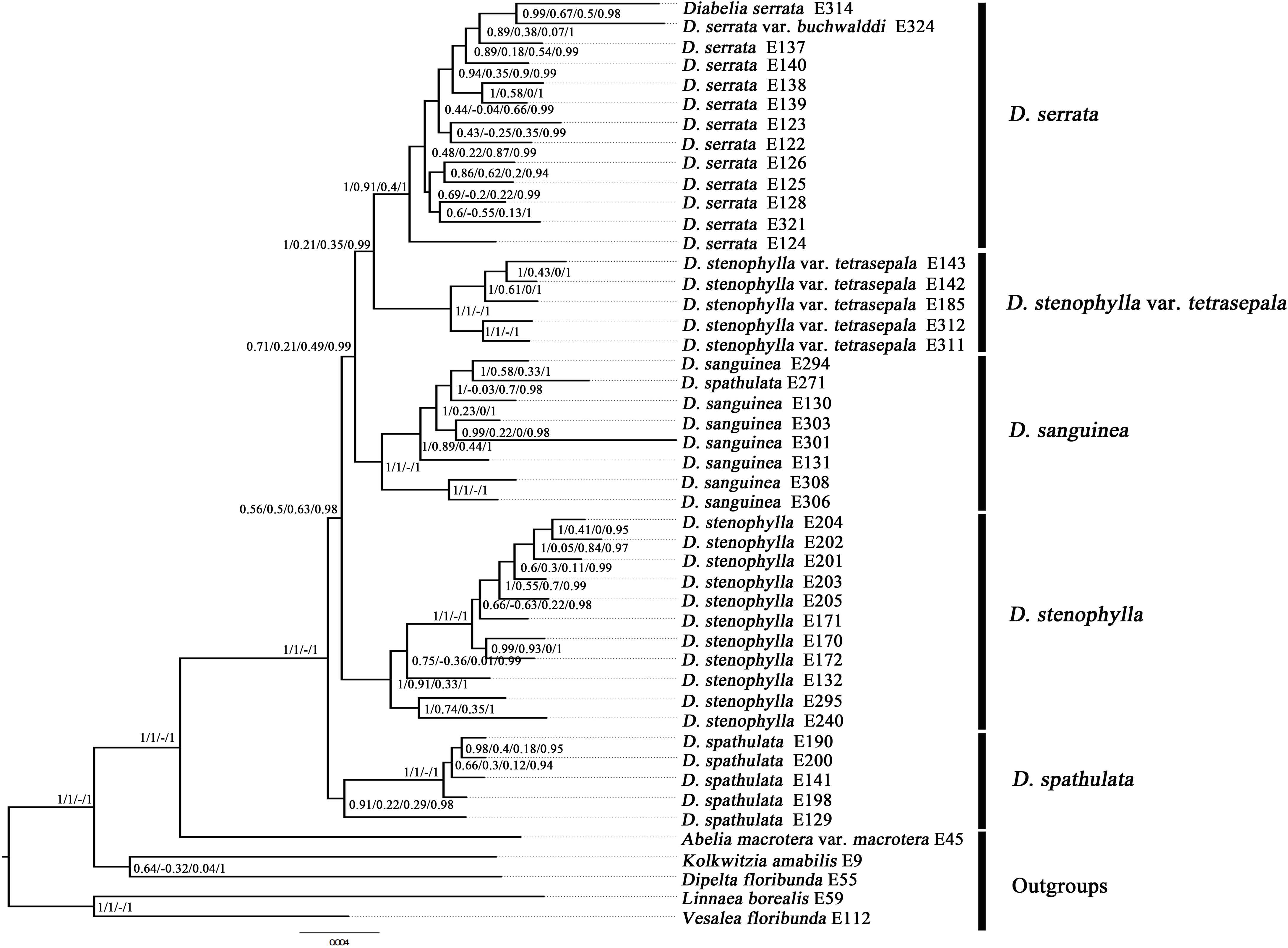
Results of the Quartet Sampling of the nuclear RAxML tree; Clade support is depicted as: Local posterior probability (LPP)/Quartet concordance (QC)/Quartet differential (QD)/Quartet informativeness (QI).

**Fig. S4.**
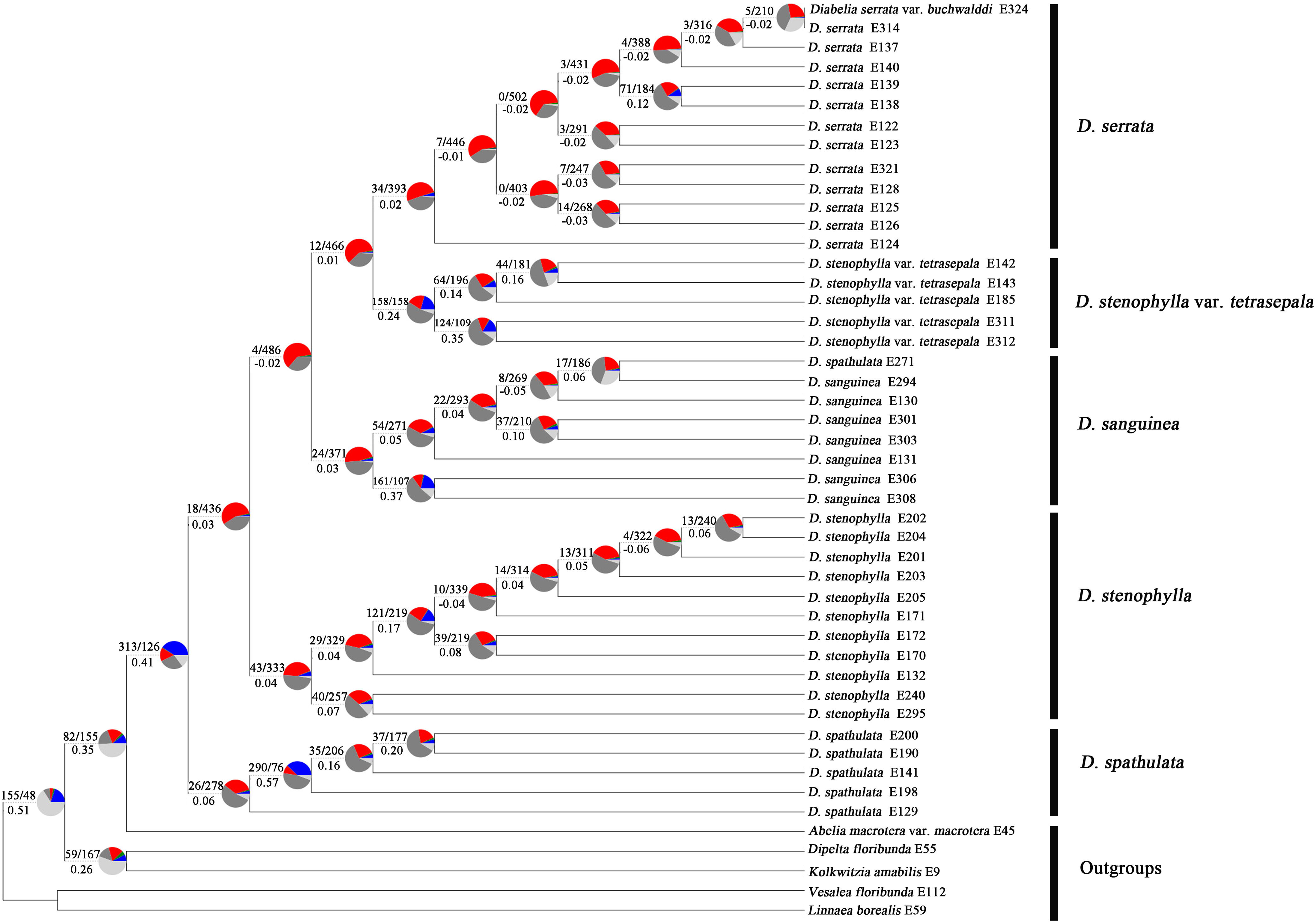
Nuclear RAxML tree. Numbers above branches indicate the number of gene trees concordant/conflicting with that node in the species tree. Numbers below the branches are the Internode Certainty All score. Pie charts next to the nodes present the proportion of gene trees that supports that clade (blue), the proportion that supports the main alternative for that clade (green), the proportion that supports the remaining alternatives (red), light gray means missing data, and dark gray means uninformative (BS < 50%).

**Fig. S5.**
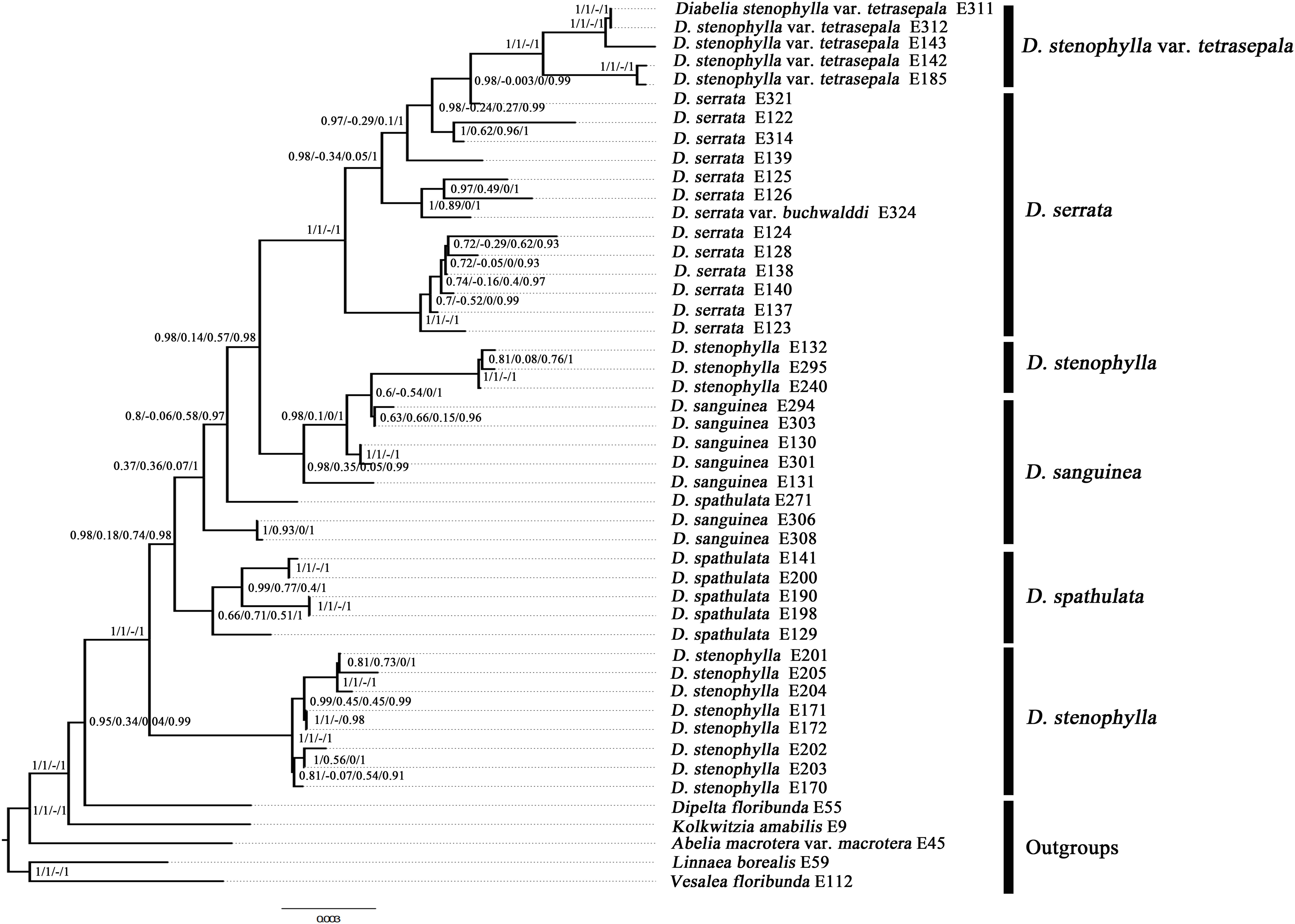
Results of the Quartet Sampling of the plastid IQ-tree tree; Clade support is depicted as: Local posterior probability (LPP)/Quartet concordance (QC)/Quartet differential (QD)/Quartet informativeness (QI).

**Fig. S6.**
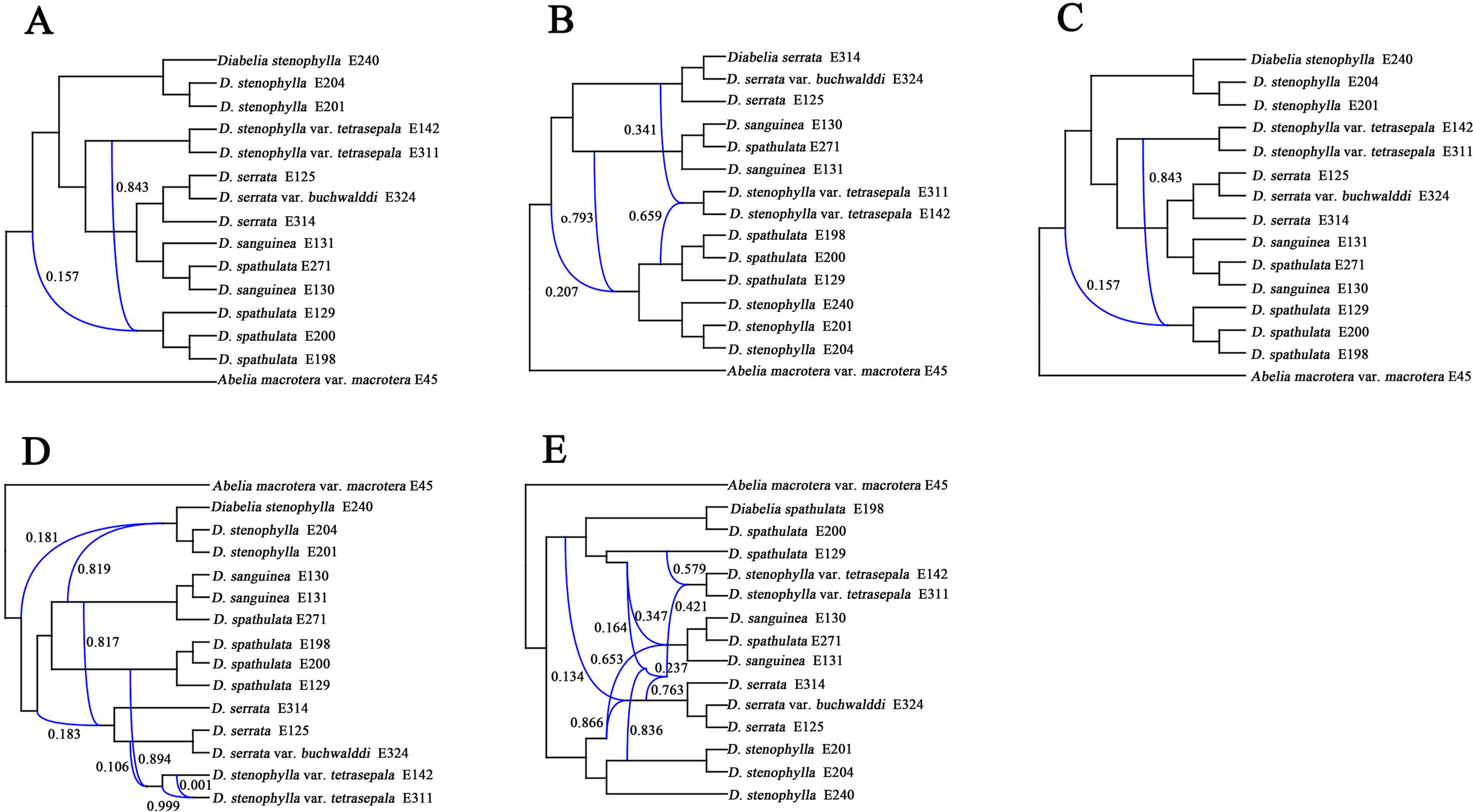
Best species networks of the selective nuclear data set estimated with PhyloNet for the 15-taxa data set. A: One hybridization event; B: Two hybridization events; C: Three hybridization events; D: Four hybridization events; F: Five hybridization events. Blue branches connect the hybrid nodes. Numbers next to blue branches indicate inheritance probabilities.

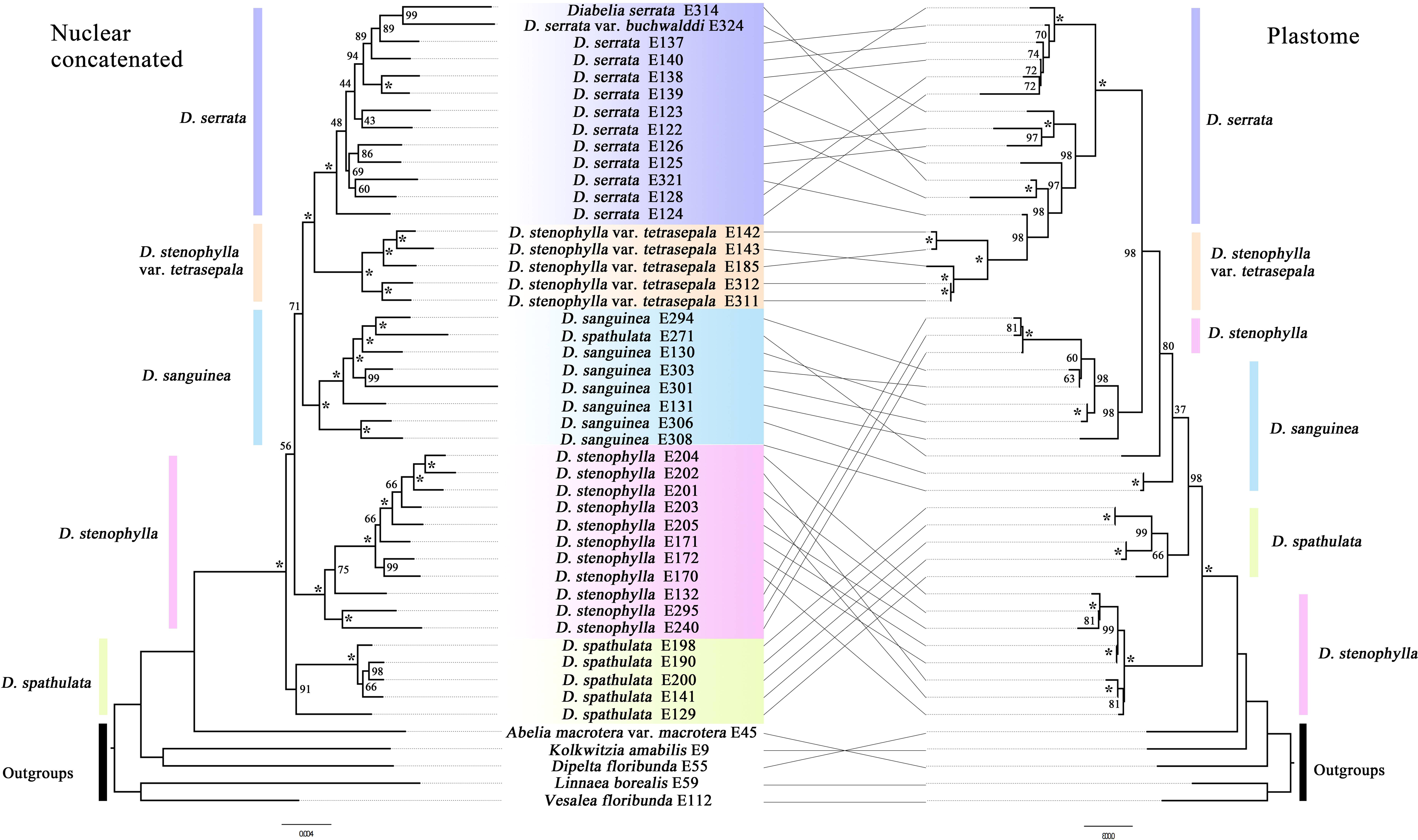

**Table S1.**
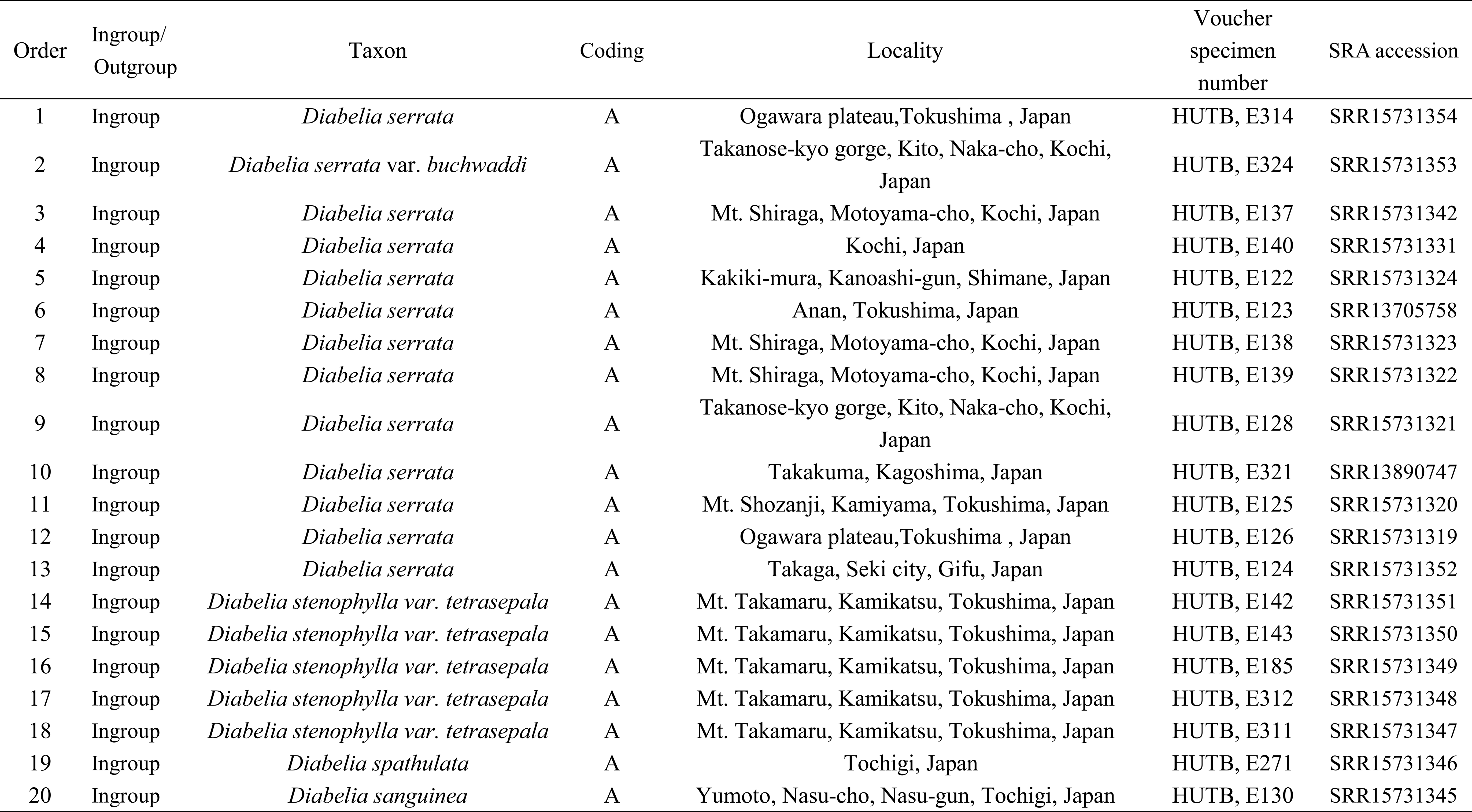
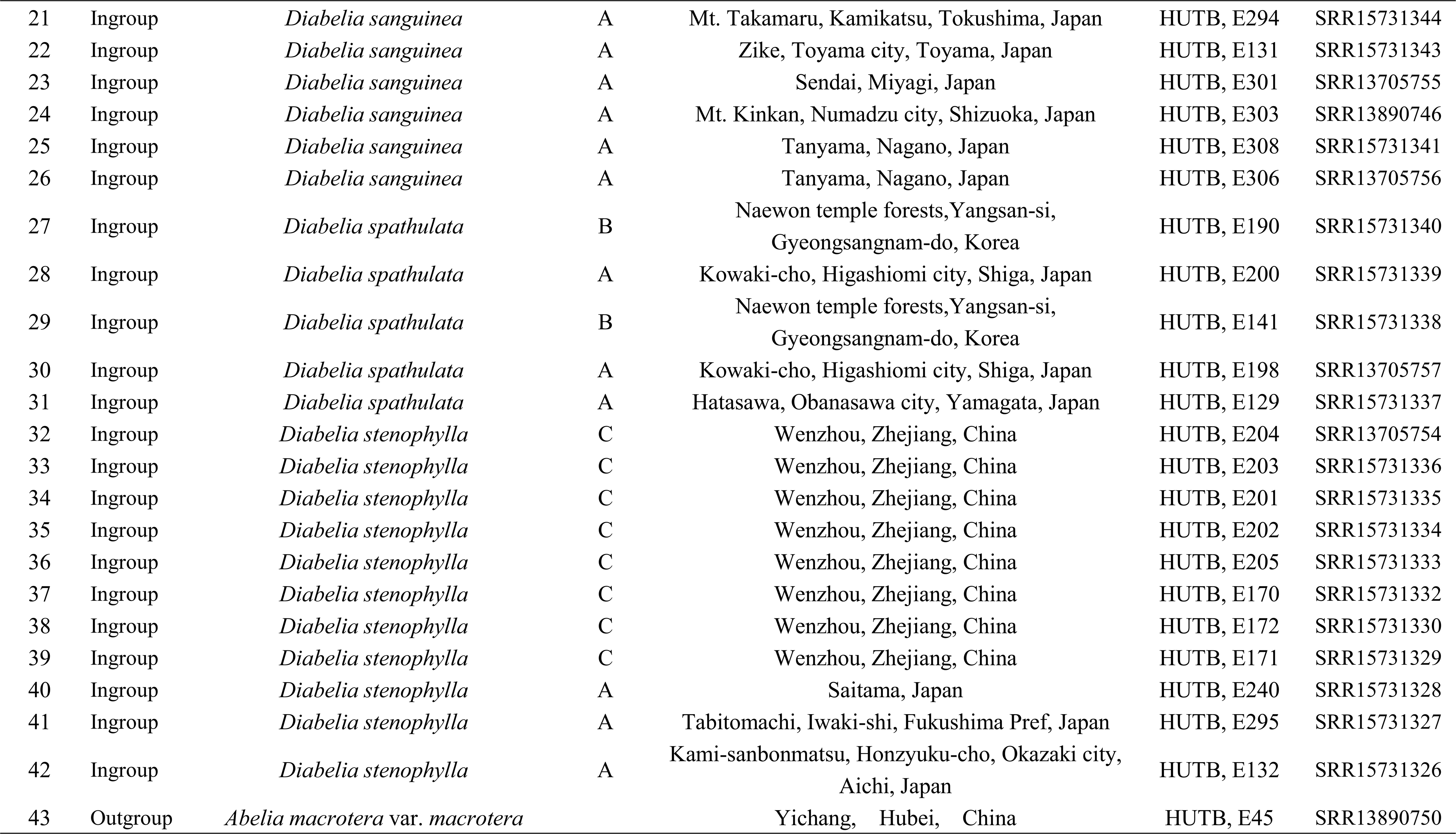
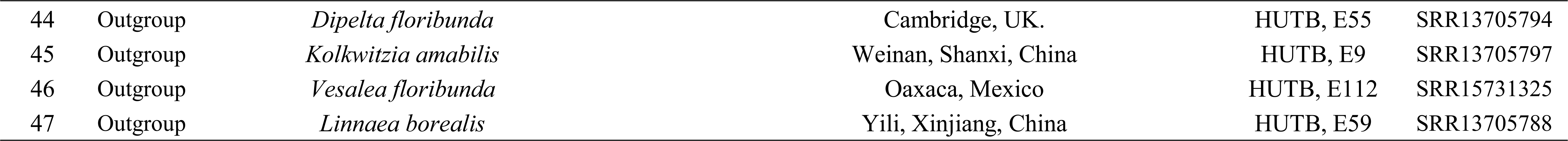
List of species and vouchers used in this study.

**Table S2.**
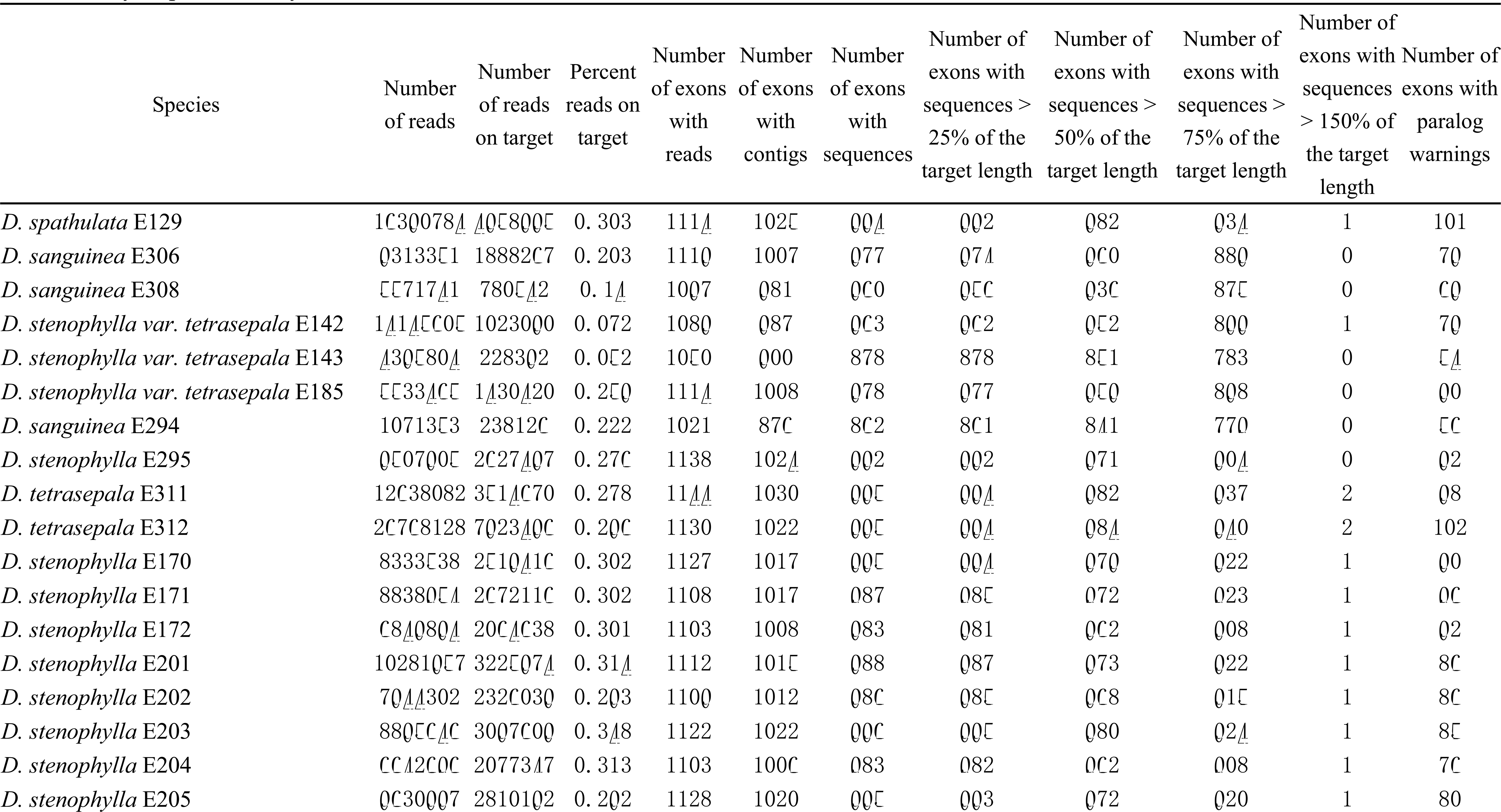
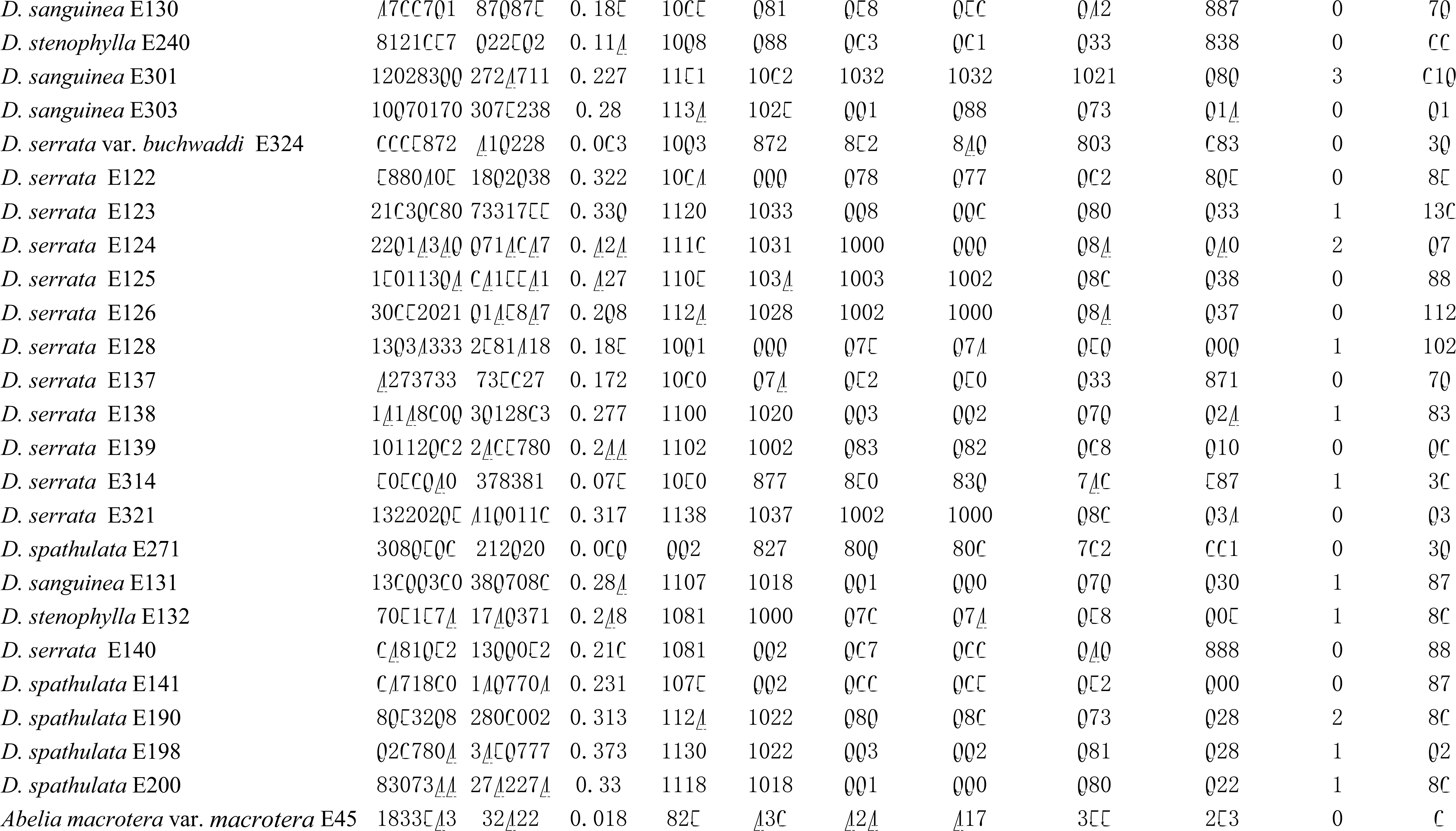
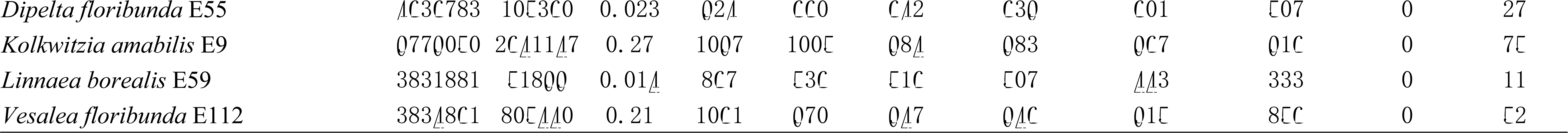
HybPiper assembly statistics.

